# Wing mechanosensory modulation of optic flow-sensitive descending neurons in butterflies

**DOI:** 10.64898/2026.01.22.701054

**Authors:** Julie Lin, Jack A. Supple, Alexandra M. Yarger, Holger G. Krapp, Huai-Ti Lin

## Abstract

Visual motion processing in flying insects is strongly modulated by behavioural state, yet the mechanisms by which mechanosensory feedback contributes to this modulation remain poorly understood. Here we show that wing mechanosensation alone is sufficient to modulate a subset of optic-flow–sensitive descending neurons (WFDNs) in butterflies. Airflow stimulation of the wings, mimicking flight conditions, increased baseline firing rates and reduced response latencies in WFDNs, without altering response gain or temporal frequency tuning. These effects indicate that mechanosensory modulation acts through mechanisms distinct from those governing other state-dependent changes in visual processing. Consistent with this interpretation, previous work has shown that baseline firing and response latency can be rapidly modulated, whereas gain changes arise through slower neuromodulatory pathways. Mechanosensory modulation was cell-type specific: the horizontally tuned WFDN_L_ neuron was consistently affected across individuals, whereas other WFDN types were largely insensitive, likely reflecting the unilateral airflow stimuli used here. Interestingly, WFDN_L_ modulation arose exclusively from mechanosensory input from the proximal area of the wing, not from distal wing deformation. This suggests that descending visual pathways require only coarse gain or excitability modulation rather than detailed information about wing shape or strain, leaving fast reflexive control of distal wing deformation to local ganglionic circuits. Together, our results demonstrate that wing mechanosensation selectively modulates visual descending pathways by altering excitability and timing rather than visual feature encoding, supporting the existence of multiple, parallel mechanisms for state-dependent modulation of descending visual signals during flight.

## Introduction

Insect flight depends on rapid closed-loop control under variable aerodynamic and inertial disturbances, requiring continuous sensory feedback about body motion and flight motor state. Because flying insects exhibit little passive stability, they rely heavily on active stabilisation supported by multimodal sensory feedback (Taylor and Krapp, 2007). This feedback includes vision (compound eyes and ocelli (Hardcastle and Krapp, 2016)), airflow sensing (antennae and wind-sensitive hairs (Budick et al., 2007)), inertial sensing (e.g., dipteran halteres (Mohren et al., 2019)), and wing-load/strain sensing (Aiello et al., 2021a; Yarger et al., 2025). Signals from these modalities are integrated to improve estimates of state changes and extend the range of perturbation frequencies over which sensory feedback remains effective (Taylor and Krapp, 2007).

Visual and mechanosensory feedback provide complementary information across an extended range of input dynamics (Hengstenberg, 1993; Schwyn et al., 2011; Sherman and Dickinson, 2004; Parsons et al., 2010). Visual motion provides rich global information about self-motion through optic flow, but its utility at high temporal frequencies is limited by delays inherent in visual processing (Taylor and Krapp, 2007). In contrast, mechanosensory pathways can provide lower-latency feedback at higher temporal frequencies, but estimates derived from these signals may accumulate bias or drift over longer timescales and can therefore benefit from slower visual correction. (Fabian et al., 2024; Verbe et al., 2020). Such complementary dynamics allow rapid mechanosensory feedback and slower, globally referenced visual feedback to act together in stabilisation reflexes, as observed across multiple insect species (Dahake et al., 2018; Hengstenberg, 1993).

A key site for multimodal integration in the insect nervous system is the population of descending neurons (DNs) that project from the brain to thoracic motor centres (Gronenberg and Strausfeld, 1990; Namiki et al., 2018). DNs sensitive to wide-field optic flow (WFDNs) coordinate stabilisation reflexes that counter unexpected perturbations to self-motion (Supple et al., 2026; Suver et al., 2016; Wertz et al., 2009a). WFDNs have been described in several insect orders, including Diptera, Orthoptera, Odonata, Hymenoptera and Lepidoptera (Ibbotson and GOODMAN, 1990; Nicholas et al., 2020; Olberg, 1981; Rowell and Reichert, 1986; Strausfeld and Bassemir, 1985; Suver et al., 2016; Supple et al., 2026) and primarily receive input from optic flow-sensitive interneurons projecting from the optic lobe (Haag et al., 2007; Strausfeld and Bassemir, 1985; Wertz et al., 2008). In flies, a population of lobula plate tangential cells (LPTCs) are matched to specific patterns of optic-flow (Krapp and Hengstenberg, 1996) and form electrical synapses with WFDNs (Haag et al., 2007; Strausfeld and Bassemir, 1985), which retain this optic flow selectivity (Wertz et al., 2009b). WFDNs are often multimodal, combining additional indicators of self-motion, such as antennal and neck movements (Olberg, 1981). In blowflies, WFDNs also combine visual information from separate sensors, the compound eyes and ocelli (Haag et al., 2007; Strausfeld and Bassemir, 1985; Wertz et al., 2008). Although less resolved than compound eyes, ocelli are faster, and their convergence onto WFDNs effectively reduces the latency of compound eye inputs (Parsons et al., 2010).

In addition to integrating information across sensory modalities, insects adjust the operating properties of individual sensory pathways according to behavioural state (Longden and Krapp, 2009; Maimon et al., 2010). For example, in flies, motion-sensitive visual circuits are strongly modulated by locomotor state (Longden et al., 2014; Maimon et al., 2010; Tuthill et al., 2014). LPTCs show increased response gain to the same visual motion stimuli during locomotion compared to stationary animals (Longden et al., 2014; Maimon et al., 2010). Locomotion also shifts LPTC frequency tuning to progressively higher image speeds, consistent with matching visual sensitivity to the faster temporal dynamics of sensory feedback during movement (Chiappe et al., 2010; Jung et al., 2011; Longden et al., 2014). These changes are also accompanied by elevated baseline membrane potentials (Maimon et al., 2010), increased firing rates (Jung et al., 2011), reduced visual response latencies, and higher information rates (Longden and Krapp, 2010, 2009).

State-dependent modulation of motion vision is linked to octopaminergic modulation (Longden and Krapp, 2010; Suver et al., 2012). In *Drosophila*, octopaminergic neurons projecting to the optic lobes are necessary and sufficient to produce locomotion-like changes in LPTC activity (Suver et al., 2012). However, octopaminergic signalling does not appear to account for all locomotion-related changes in visual processing. In *Drosophila* LPTCs, changes in baseline membrane potential are not reproduced by octopaminergic application (Suver et al., 2012), and occur on a much faster timescale than other octopamine-related changes in visual gain (Maimon et al., 2010). Furthermore, nutritional deprivation counteracts locomotion-related modulation of motion vision to conserve energy, although locomotion-related latency decrements remain unaffected (Longden et al., 2014). Together, these findings indicate that state-dependent modulation of motion vision is implemented through multiple mechanisms.

Convergence of signals from different sensory modalities onto shared neural circuits may provide an additional mechanism for state-dependent modulation of sensory processing (Rimniceanu et al., 2023; Santer et al., 2006; Wasserman et al., 2015). Past work has described multi-modal integration in the context of improving the accuracy and precision of sensory estimates (Olberg, 1981; Parsons et al., 2010) but less is known about whether signals from one modality can convey information about locomotor state changes and thereby modulate processing in another sensory pathway. Ascending neurons (ANs) convey information from thoracic ganglia to the brain and are thought to transmit proprioceptive and motor related information to cephalic sensory circuits (Chapman et al., 2018; Chen et al., 2023; Cheong et al., 2024; Fujiwara et al., 2022). ANs may be involved in activating octopamine signalling (Suver et al., 2012). They could also provide faster state-related signals to visual circuits, although their contribution to changes in baseline activity or response latency remains unresolved. Alternatively, descending neurons themselves could provide a site at which thoracic mechanosensory signals directly converge with visual input from the brain, allowing rapid modulation of descending responses without first retuning upstream visual circuits.

Wing mechanosensory feedback provides one potential source of the rapid thoracic sensory input proposed above, because insect wings experience substantial aerodynamic loading and deformation during flight. Insect wings are therefore not only aerodynamic actuators but also sensory surfaces containing distributed mechanoreceptor arrays. Whole-wing mapping across insects has revealed spatially organised arrays of mechanosensors distributed across the wing surface and veins (Fabian et al., 2022; Stanchak et al., 2024; Yarger et al., 2025; Yoshida and Emoto, 2011). Across insects, campaniform sensilla (CS) transduce local cuticular strain (Aiello et al., 2021b), and their placement across the wing constrains the mechanical inputs they encod (Stanchak et al., 2024).

In this study we tested whether wing mechanosensory input evoked by externally applied airflow modulates optic-flow-sensitive descending neurons in butterflies. Butterflies have relatively large wings and low wingbeat frequencies compared with many other flying insects, with wingbeat frequencies of approximately 10–15 Hz reported in previous studies (Fei and Yang, 2016; Johansson and Henningsson, 2021; Srygley and Thomas, 2002). Butterflies frequently engage in gliding as well as flapping flight (Harris et al., 1999; Stylman et al., 2020), such that airflow across the wings in the absence of active flapping may be a frequently encountered feature relevant for flight control. We therefore investigated whether mechanosensory input generated by airflow applied to the wings is sufficient to modulate WFDN activity in the absence of active wing flapping.

## Results

### Wing airflow modulates spike rates for specific WFDN cell types

To investigate whether wing mechanosensory input modulates visual descending responses, we measured extracellular action potentials from the cervical connective and presented moving gratings in the frontal visual field with and without airflow across the wings (Figure 1A-B). The wings were immobilised in a gliding-like posture, with the leading edge approximately perpendicular to the body axis (Figure 1A). The antennae were removed and their bases covered with beeswax to minimise potential antennal mechanosensory inputs to DNs. We recorded four WFDN response classes: WFDN_L_ (9 cells from 9 animals), WFDN_M_, (4 cells from 4 animals), WFDN_V_ (4 cells from 4 animals), and WFDN_D_ (4 cells from 4 animals). WFDN_L_ and WFDN_M_ responded preferentially to horizontal motion in opposite directions, whereas WFDN_D_ and WFDN_V_ responded preferentially to ventral-to-dorsal and dorsal-to-ventral motion, respectively (Figures 1C and 2A–D; (Supple et al., 2026)).

**Figure 1:**
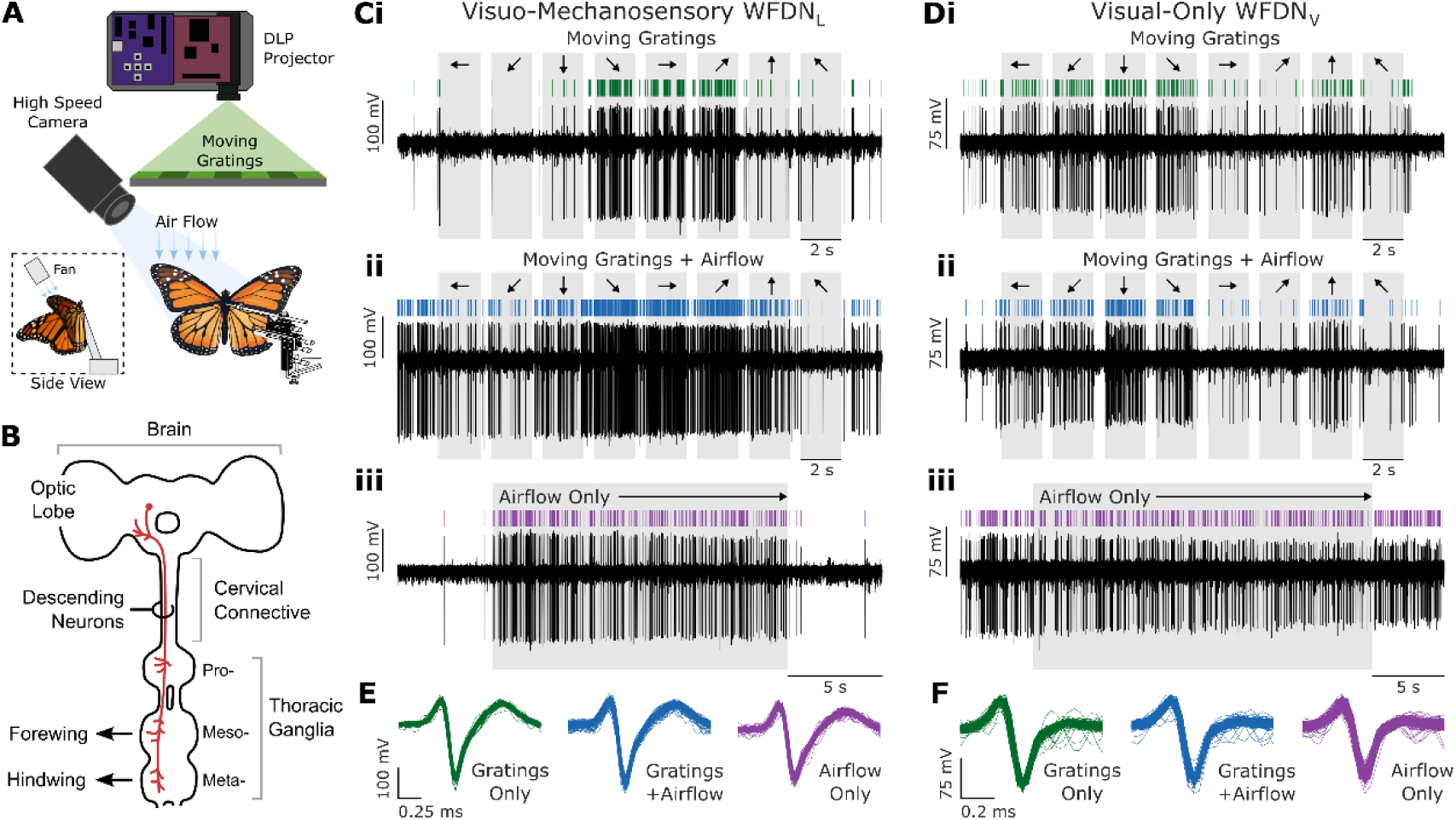
Butterfly wide-field optic flow-sensitive descending neuron (WFDN) responses to visual stimuli and aeroelastic wing deformation. **(A)** Experimental setup. Butterflies were mounted ventral-side up with a DLP projected image positioned in the frontal visual field. Animals were presented with gratings moving sequentially along different directions with and without airflow directed towards the butterflies’ right forewing, with an angle of attack of 0°. Aeroelastic wing flutters were filmed at 868.8 fps. **(B)** Schematic of the butterfly visuomotor descending-neuron pathway, viewed from the ventral aspect of the animal, such that the animal’s right is on the left-hand side. Extracellular measurements of butterfly descending neurons (DNs, red) were recorded from the animal’s right cervical connective. **(C)** Example visuo-mechanosensory descending neuron WFDN_L_ sensitive to both gratings motion and airflow across the wing. (i) Response to gratings moving along eight directions in the frontal visual field without airflow. Arrows indicate direction of grating motion in the animal’s frame of reference, i.e., animal’s left, down (ventral), right, and up (dorsal). (ii) Response to the same moving gratings with airflow across the wings. (iii) Response to airflow across the wings, without any visual stimulation. **(D)** Example visual-only descending neuron WFDN_V_ that did not exhibit notable response to airflow. Note that both example neurons in C-D were from the same animal. Panels (i-iii) same as (C). **(E-F)** Spike sorted units from each stimulus combination in the example traces (C-D).

**Figure 2:**
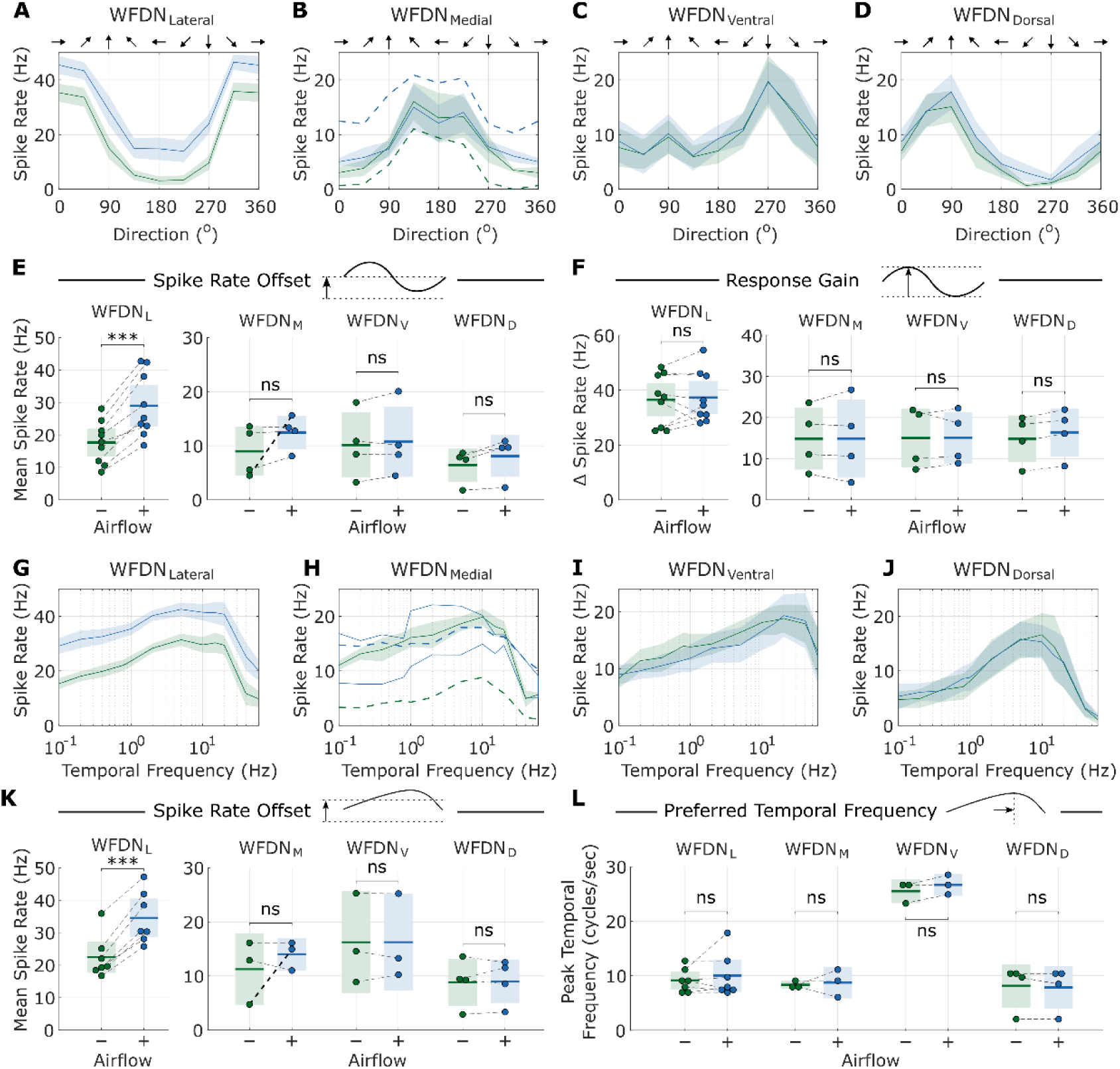
Aeroelastic wing flutter modulates spike rates for selected WFDN cell types. **(A-D)** Directional tuning curves for four commonly encountered WFDN cell types sensitive to different directions of local motion in the frontal visual field. (A) WFDN_L_ : N=9 cells from 9 animals; (B) WFDN_M_ : N=4 cells from 4 animals; (C) WFDN_V_ : N=4 cells from 4 animals. (D) WFDN_D_ : N=4 cells from 4 animals. Response to visual motion without airflow are in green, and vision+airflow in blue. Whilst DNs sensitive to 0° gratings motion (B) tended not to change response when airflow was presented, one animal displayed a substantial increase in spike rate when presented with airflow (indicated by dashed curves). Arrows indicate grating direction in animal’s frame of reference: 0° left, 90° dorsal, 180° right, 270° ventral. **(E-F)** Statistical analysis of data in panels A-D **(G-H)** Descending neuron temporal frequency responses for gratings moving along the preferred direction in the frontal visual field. Response to visual motion without airflow are in blue, and vision+airflow in red. Only two frequency tuning curves were measured under airflow conditions for DNs sensitive to 0° gratings motion (H). Individual tuning curves are shown in solid red lines. An average of three tuning curves is shown for vision- only conditions (blue). Dashed blue/red curves show the response of the airflow-sensitive 0° DN, corresponding to the same neuron in (B). **(K-L)** Statistical analysis of data in panels G-H.

WFDN_L_ showed a statistically significant increase in mean spike rate when visual motion was presented together with airflow applied to the wings (paired t-test; Bonferroni-corrected p = 5×10⁻⁴; N=9 animals). WFDN_L_ mean spike rate increased by 11.3±4.5 spikes/s, from 17.6±6.5 to 29.0±9.7 spikes/s (mean±std; Figure 2E). Whilst WFDN_D_ showed a subtle increase in mean spike rate from 6.4±3.1 to 8.1±3.9 spikes/sec, this was statistically insignificant (paired t-test; Bonferroni-corrected p=0.47, N=4 animals). There was no consistent change in mean spike rates for WFDN_M_/WFDN_V_ visual responses with airflow (WFDN_M_ : 9.0±4.6 to 12.4±3.2, and WFDN_V_ : 10.1±6.1 to 10.7±6.6 spikes/sec for visual responses without and with airflow, respectively). Notably, one WFDN_M_ neuron did exhibit a substantial increase in spike rate with airflow akin to WFDN_L_ ; however, this was only observed once and was therefore not interpreted as a consistent WFDN_L_ response (dashed curve, Figure 2B). Furthermore, differences in WFDN airflow sensitivity were also observed within individuals, such that modulations in WFDN_L_ were seen alongside airflow-insensitive WFDN_V_ within the same preparation (Figure 1C-D). This within-animal difference indicates that airflow sensitivity is not a general feature of WFDNs, but is specific to particular WFDN classes.

### Wing airflow does not alter optic flow processing

Locomotor state can alter both visual response gain and temporal-frequency tuning in insect motion-sensitive neurons (Maimon et al., 2010; Suver et al., 2012; Longden et al., 2014). We therefore tested whether airflow-evoked modulation of WFDNs altered these visual response properties by comparing directional response gain (Figure 2F) and temporal-frequency tuning (Figure 2G–L) between airflow-off and airflow-on conditions.

There was no statistically significant difference in visual directional response gain for any WFDN type after correcting for multiple comparisons (paired t-test; Bonferroni-corrected p>0.05; Figure 2F). WFDN_D_ showed the largest increase in directional response gain, from 14.9 ± 5.8 spikes/s without airflow to 16.4 ± 6.0 spikes/s with airflow (paired t-test, unadjusted p = 0.007). However, this effect did not remain significant after Bonferroni correction (adjusted p = 0.06; Figure 2F). No WFDN class showed a significant change in directional response gain after correction for multiple comparisons.

To test whether mechanosensory modulation altered the dynamics of WFDN visual responses, we compared temporal-frequency tuning between airflow-off and airflow-on conditions. Across the temporal-frequency series (Figure 2F), visual temporal frequency tuning showed a statistically significant baseline shift in average spike rates across a range of temporal frequencies for visuo-mechanosensory WFDN_L_ (Figure 2K; mean WFDN_L_ spike rate: 22.4±6.5 and 34.6±8.0, without/with airflow, respectively; Bonferroni-corrected p=4.3×10^-4^). Other WFDN types did not display any statistically significant difference in average spike rate offset. The same WFDN_M_ recording highlighted above showed a large airflow-associated increase in mean firing, from 4.7 to 15.0 spikes/sec (without/with airflow; dashed line, Figure 2H), but this response was not consistent across WFDN_L_ recordings.

There was no statistically significant difference in the temporal frequency producing the maximum spike rates between airflow and no-airflow conditions, for any WFDN type (paired t-test; Bonferroni-corrected p>0.5; Figure 2L). Whilst airflow increased the baseline of WFDN_L_ responses, the preferred temporal frequency was unchanged (9.1±2.2 and 10.0±4.0 cycles/sec, without/with airflow, respectively). Preferred temporal frequencies also differed across WFDN classes: WFDN_L_, WFDN_M_ and WFDN_D_ preferred frequencies around 8–10 cycles/s in the absence of airflow (9.1 ± 2.2, 8.8 ± 2.5, and 7.9 ± 4.0 cycles/s, respectively), whereas WFDN_L_ showed a substantially higher preferred temporal frequency of 26.8 ± 1.8 cycles/s (Figure 2L). Together, these results show that airflow increased WFDN_L_ spike rates without producing a detectable shift in directional response gain or preferred temporal frequency.

### Wing airflow-induced spike rate modulation reduces visual latency

We next investigated whether airflow-induced increases in baseline firing affect visual response latency (Figure 3). We hypothesised that an increased synaptic drive from wing afferents shifts WFDN membrane potential closer to spike threshold, such that visual inputs require less time to elicit action potentials and therefore reducing response latency.

**Figure 3:**
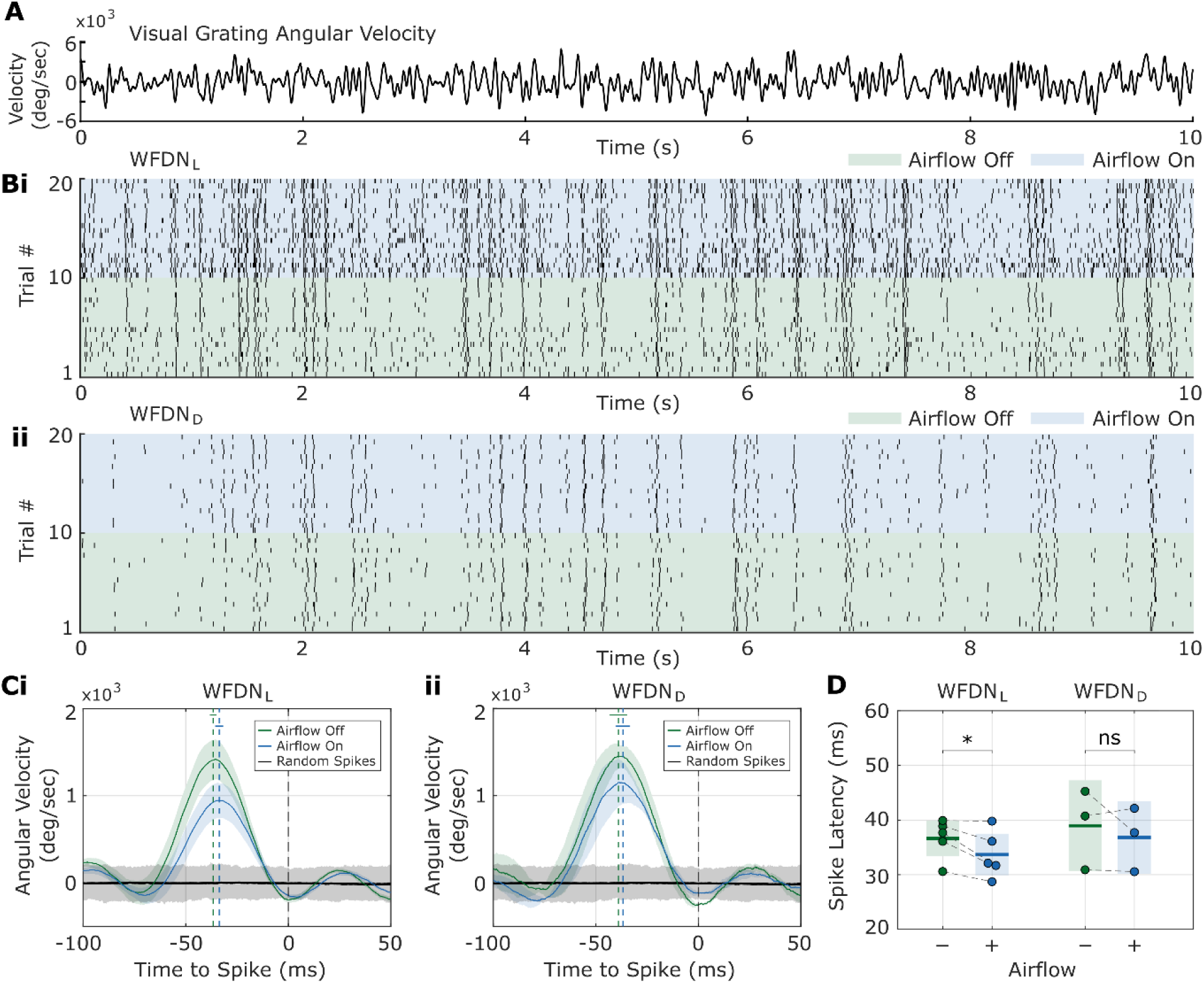
Wing airflow-induced spike rate modulation reduces visual latency. **(A)** Gratings moved along the preferred direction with band-limited Gaussian-noise velocity. **(B)** Example spike raster for (i) WFDN_L_ and (ii) WFDN_D_ responding to repeated presentations of the gratings stimulus in (A) without (green) and with (blue) airflow across the ipsilateral wing. **(C)** Spike-triggered averages (STAs) of grating velocity without (green) and with (blue) airflow for (i) WFDN_L_ (N=5 animals) and (ii) WFDN_D_ (N=3 animals). Curves show mean±SEM. Spike time is at 0 ms (vertical dashed black line). Vertical dashed green and blue lines show average latency without and with airflow, respectively. Small intersecting horizontal lines show latency SEM. Black line and grey boundary shows STA mean and 95 percentiles for randomised spike times. **(D)** Spike latency measurements corresponding to the temporal offset of the STA grating velocity peak preceding the spike time. Lines and shaded regions show mean±SEM, circles show individual animals. Paired t-test WFDN_L_ p=0.04; WFDN_V_ p=0.51.

To estimate visual response latency, we calculated the spike-triggered average (STA) of grating velocity, representing the average visual velocity profile preceding a spike, and defined latency as the interval between the preceding STA velocity peak and spike time (Figure 3C–D). For WFDM_­_, the latency between the peak grating velocity and spike time was 36.6±3.7 ms without airflow and 33.7±4.3 ms with airflow, a statistically significant decrease of 2.9±2.2 ms (i.e. -7.9%, paired t-test; p=0.04; N=5 animals; Figure 3D). In contrast, WFDN_V_ had no difference in response latency with and without airflow (without airflow: 39.0±7.3 ms; with airflow: 36.8±5.9 ms; paired t-test p=0.51; Figure 3D).

### Mechanosensory input to WFDN_L_ derives from the wing base

We next investigated the spatial and frequency-dependent properties of wing mechanosensory input to WFDN_L_ (Figure 4). We first used localised, frequency-modulated sinusoidal airflow to identify the wing region and airflow frequency that elicited the strongest responses (Figure 4A-B). WFDN_L_ showed no response to stimulation in the distal forewing (Figure 4Bi, pink) but was tuned to ∼60 Hz stimulation at the forewing base (full width at half maximum: 35-105 Hz; N=2 animals; Figure 4Bi, blue). In contrast, mechanosensitive-only interneurons recorded from the meso-thoracic ganglion responded to stimulation in the distal forewing (Figure Bii, pink) with a preferred frequency of 80 Hz (full width at half maximum: 30-190 Hz; N=2 animals).

**Figure 4:**
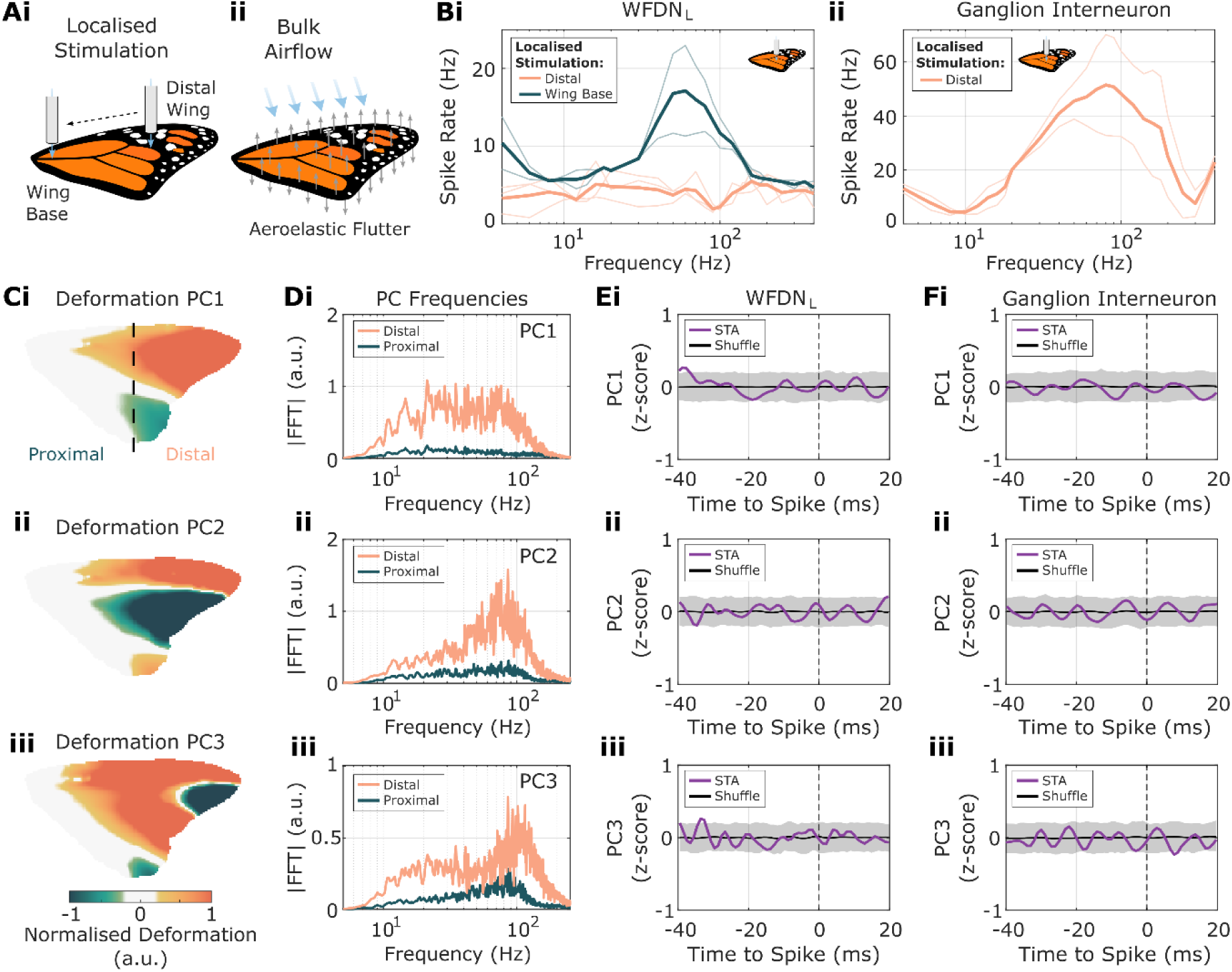
Mechanosensory input to WFDN_L_ originates from the wing base. **(A)** Mechanosensory stimuli consisted of either (i) localised stimulation with sinusoidally modulated air flow, or (ii) bulk airflow across the wing (as used in Figures 1-3). Grey arrows indicate aeroelastic wing deformation. Localised stimulation is applied ventrally, normal to the wing surface; bulk airflow is applied with an angle-of-attack of 0°. **(B)** Mechanical frequency sensitivity tuning curves measured from localised sinusoidal airflow stimulation for (i) WFDN_L_ and (ii) non-visual mechanosensitive interneurons in the meso-thoracic ganglion. Localised stimuli were tested at the distal wing (pink) and wing base (blue). Thick likes show averages, thin lines show individual cells. WFDN_L_ : distal wing N=3 animals, wing base N=2 animals; ganglion interneuron: N=2 animals. **(C)** The first three principal components (PCs) of aeroelastic wing deformation under bulk airflow. Delineation of distal and proximal wing regions indicated by dashed line in (i). **(D)** Fourier transform of wing deformation PCs in (C). **(E)** WFDN_L_ spike triggered average (STA, purple line) of wing deformation PCs 1-3. Spike time is at 0 ms (vertical dashed black line). Black line and grey boundary show STA and 95 percentiles for randomised spike times. N=1 cell, corresponding to one cell from (Bi). **(F)** Mechanosensitive ganglion interneuron STA (purple line) of wing deformation PCs 1-3. Same format as (E). N=1 cell, corresponding to one cell from (Bii).

To further explore how neuronal mechanosensitivity correlated with the aeroelastic wing deformation under the bulk airflow used in our previous experiments (Figures 1-3), we investigated the relationship between neural responses and steady-state wing deformation from high-speed videos (Figure 4C-F). We first decomposed wing deformation using principal component analysis (PCA; Figure 4C). The first 3 principal components (PCs) captured ∼90% of wing deformation variance, with most movement occurring at the distal forewing (Figure 4C-D). PC1 deformation contained power across a broad range of frequencies until attenuating above 80 Hz (Figure 4Di), whilst PC2-3 deformation was focused at 80 Hz and 100 Hz, respectively (Figure 4Dii-iii). We next calculated STAs of wing deformation PCs 1-3 for WFDN_L_ and ganglion interneurons (Figure 4E-F). In contrast to our localised airflow stimuli, we did not observe any clear correlation between spike timing and these three highest PCs, for either WFDNL or the ganglion interneuron (Figure 4E-F). Together, these results suggest that WFDN_L_ mechanosensitivity is preferentially associated with proximal-wing stimulation, while the dominant wing-deformation PCs showed no clear spike-triggered relationship in the cells analysed.

## Discussion

State-dependent modulation of neural processing reflects a general computational strategy whereby the system-level dynamic range is extended by shifting the relatively narrow operation range of individual neurons to meet context-dependent requirements. In Diptera, changes in internal state - such as locomotion activity or satiation status - modulate optic flow processing in both graded (Maimon et al., 2010) and spiking (K. D. Longden et al., 2014) LPTCs. Locomotion-related modulation produces a variety of changes that shift visual processing toward a faster operating regime, consistent with the increased temporal dynamics of sensory feedback during movement (Maimon et al., 2010). These include increased visual gain (Maimon et al., 2010), a shift in temporal-frequency tuning toward higher image velocities (Jung et al., 2011), elevated baseline membrane potential (Maimon et al., 2010) and firing rates (K. D. Longden et al., 2014), shortened response latencies, and increased information rates (Longden and Krapp, 2010).

Our results align with these previous findings by demonstrating modulation of optic flow-sensitive DNs during mechanosensation of airflow across the wing, which would occur during flight (Figures 1-3). However, because the wings were immobilised in a gliding-like posture, these experiments isolate mechanosensory responses to airflow across stationary wings and do not capture the additional mechanosensory signals generated during active wing flapping. Notably, mechanosensory-induced modulation was limited to increased baseline firing rates (Figure 2E,K) and shortened response latencies (Figure 3) for WFDN_L_. It did not change other properties of visual motion detection such as directional preference, response gain (Figure 2F) or temporal frequency tuning (Figure 2L). This suggests that latency modulation arises from separate mechanisms to the modulation of other properties of optic flow processing. Indeed, in *Drosophila*, changes in baseline membrane potential are rapid, closely following flight state (Maimon et al., 2010), whilst flight-induced elevation of visual gain has slower dynamics arising from octopamine neuromodulation (Suver et al., 2012). Similarly, nutritional deprivation elicits energy-conserving constraints on locomotion-related modulation, including cancellation of increased baseline firing rates and response gain, whilst locomotion-related latency decrements remain unaffected (Longden et al., 2014). Our finding that wing mechanosensation effects baseline firing and response latency, but not response gain or temporal frequency tuning, further suggests that multiple mechanisms underly state-dependent changes in visual processing.

Motion detection first occurs in optic lobe T4/T5 cells, which receive dendritic inputs from the medulla/lobula, respectively (Fischbach and Dittrich, 1989; Maisak et al., 2013). Visual gain modulation likely depends on a range of mechanisms including changes in presynaptic excitation and inhibition (Longden and Krapp, 2010; Tuthill et al., 2014). Temporal frequency tuning is determined primarily by the delay time constants of elementary movement detection (Harris et al., 1999; Jung et al., 2011; Strother et al., 2018). Changes in optic flow processing therefore requires modulating these upstream neurons located in the optic lobe (Strother et al., 2018; Tuthill et al., 2014). In contrast, elevated baseline firing and latency reduction can both be achieved by increasing the resting membrane potential of a given neuron closer to spike threshold (Maimon et al., 2010; Santer et al., 2006; Tuthill et al., 2014). Whilst ascending mechanosensory information has been shown to modulate LPTCs (Cheong et al., 2024; Fujiwara et al., 2022, 2017), DN subthreshold membrane potential could be modulated directly by synaptic input within the thoracic ganglia.

Our results show that mechanosensory modulation differs among WFDN cell types (Figure 2). WFDN_L_, which responds preferentially to horizontal wide-field visual motion towards the ipsilateral side (with respect to the cervical connective), was consistently modulated by airflow stimulation across animals. All other WFDN types were not affected by airflow, except one instance of WFDN_L_, which responds preferentially to horizontal wide-field visual motion towards the contralateral side (Figure 2B,H). The restriction of modulation to horizontal WFDN types could reflect our unilateral airflow configuration, in which only the animal’s right wings were stimulated. This lateralised asymmetry could explain the absence of modulation in vertical WFDN types, which may instead require vertical asymmetries. Indeed, airflow was oriented at an angle of attack of 0° relative to the wing leading edge, producing minimal asymmetry along the vertical axis. If vertical WFDN types are preferentially modulated by vertical airflow components, we predict similar modulation analogous to WFDN_L_ under non-zero angles of attack.

Alternatively, vertical WFDN types may remain unmodulated regardless of airflow orientation, which could reflect differences in the function of horizontal and vertical WFDN types during normal flight control. Butterflies experience substantial ∼10 Hz pitching oscillations with each wing stroke in flight (Fei and Yang, 2016), and vertical WFDNs may function to stabilise these oscillations. Indeed, the difference in temporal frequency maxima for WFDN_V_ and WFDN_D_ (Figure 2F) may reflect tuning to the different pitching velocities experienced throughout the wing stroke cycle. Because vertical visual motion generated by oscillatory pitching movements and wing mechanosensory feedback vary periodically across wing strokes, additional airflow-induced modulation of vertical WFDNs may introduce response variability that disrupts periodic pitch control. Interestingly, differences between vertical and horizontal motion detection have also been reported in Diptera, with horizontal LPTCs more strongly tuned for observability, whereas vertical LPTCs are biased towards controllability and disturbance sensitivity (Humbert et al., 2026).

What is the function of wing mechanosensation-induced modulation of WFDN_L_ activity? Spike latency is reduced by ∼8%, comparable to the ∼6-13% reduction induced by octopamine in flies (Longden and Krapp, 2010, 2009), presenting a notable increase in the dynamic performance of optomotor reflexes. Increased spike rates may in turn increase the reliability of descending control of motor circuits, potentially increasing the probability of muscle contraction for a given descending signal. The preferred mechanosensory frequency of ∼60 Hz for WFDN_L_ suggests that wing mechanosensation is not tuned to the ∼10 Hz wing stroke cycle. In fact, aeroelastic wing deformation power declines above ∼80–100 Hz across all PCs (Figure 4D). Whilst PC1 exhibits broadband deformation power spanning 10-100 Hz (Figure 4Di), higher-order PCs are concentrated towards 80-100 Hz (Figure 4Dii-iii). The correspondence between aeroelastic wing deformation spectral power and WFDN_L_ mechanosensitive frequency tuning is consistent with a role in monitoring high-frequency deformation indicative of wing loading and/or aerodynamic perturbations, rather than the wing stroke cycle. WFDN_L_ was only modulated by mechanosensory input from the wing base, rather than from the distal wing. This suggests that descending visual signals only require gain modulation without detailed information on the actual wing deformation. This leaves the fast reflexive circuits within the ganglion, where we found sensory correspondence to distal deformation of the wing, key to more intricate wing deformation-related control.

Overall, our findings demonstrate that wing mechanosensation alone can modulate selected WFDN cell types in butterflies. This modulation does not impact optic flow processing, instead increasing baseline firing rates and reducing spike latency. Future work is required to study whether this modulation depends on airflow asymmetries. In addition, further analysis of butterfly wing deformation dynamics during aerodynamic perturbations may provide additional insight into the functional significance of WFDN modulation. Mechanosensory input may therefore provide a state-dependent signal that adjusts the excitability and response latency of WFDNs according to the mechanical state of the wing. Such modulation could allow visually detected changes in body motion to be transmitted more rapidly or reliably to the flight motor system when the wings are aerodynamically loaded. Whether this modulation arises through direct mechanosensory input within the thoracic ganglia or indirectly through ascending pathways remains to be determined.

## Methods

### Animals

Experiments were performed on adult Monarch butterflies (*Danaus plexippus*) obtained from a commercial supplier (The Entomologist Ltd, UK). Butterflies were delivered as pupae and reared to eclosion at 25°C, 60% humidity on a 12/12hr light/dark cycle. Adults were fed sugar solution (10% honey in water) daily. Experimental individuals were used within 5-7 days post-eclosion.

### Dissection and Electrophysiology

Individuals were immobilised by chilling on ice for five to ten minutes until all voluntary movement ceased. Wings were positioned so that the axis from forewing tip to wing hinge was perpendicular to the body axis, and secured with superglue to prevent voluntary wing movements. Each butterfly was mounted ventral side up in a custom 3D-printed holder using wax that stabilised the body while providing access to the thorax. The head was restrained with fine pins and wax bridges. The proboscis, palps, legs and antennae were removed and heavily waxed to prevent additional sensory input from these structures. The ventral nerve cord was exposed ventrally. During experiments, Lepidoptera saline (in mM: NaCl 147.0, KCl 1.3, NaHCO_­_10.0, CaCL_­_4.0, pH 7.3; (Kinoshita et al., 2015)) was applied to prevent dehydration.

Extracellular recordings were obtained using sharp 2-3 MOhm tungsten microelectrodes (Model: tungsten / 125um / 15–18um / unplated /220-P02 / 15mm; Microelectrodes Ltd., UK) inserted into the animal’s right VNC. In butterflies, the mesothoracic and metathoracic ganglia are partially fused, forming a bilobed oval structure with a visible intersegmental boundary (Niven et al., 2008) (Figure 1B). Ganglion interneuron recordings targeted the mesothoracic portion that receives forewing mechanosensory input. A fine hook was placed beneath the ganglion to elevate and stabilise the tissue and to serve as the ground reference electrode. Descending neurons were recorded from the cervical connective as described previously (Supple et al., 2026).

Extracellular action potentials were amplified and digitised via an RHD 16-channel bipolar-input headstage (Intan, USA) using the Open Ephys acquisition system at 30 kHz and monitored in real time with the Open Ephys GUI.

### Experimental Setup and Stimuli

Visual stimuli were delivered to a rear projection screen in the insect’s frontal visual field, using a LightCrafter DLP3010 (Texas Instruments, USA) at 360 frames per second (fps). A 16×9 cm projection screen was positioned 20 cm from the animal, subtending a visual angle of 44°x25°. Visual patterns were generated in MATLAB using Psychtoolbox-3 (Brainard, 1997; Pelli, 1997). Square-wave gratings were used for direction tuning (0-315° in 45° steps, 2 s per direction; temporal frequency = 5 Hz; spatial wavelength = 10°/cycle) and spatiotemporal tuning at a fixed preferred direction (5 s; spatial wavelength = 10°/cycle; temporal frequencies = 0.1-60 Hz). For response-latency measurements, sine-wave gratings moved along each neuron’s preferred direction axis according to a 0.1–20 Hz band-limited Gaussian-noise velocity profile for 10 s. The same velocity sequence was repeated for a minimum of 10 trials under both airflow-off and airflow-on conditions. Visual stimuli timings were synchronised with electrophysiology by recording a blinking trackbox in the corner of the projected image on the electrophysiology AUX channels.

Airflow stimuli consisted of either bulk airflow from the fan, or localised sinusoidal airflow oscillations. Bulk airflow was produced by a fan positioned ∼5 cm from the forewing at an angle-of-attack of 0° relative to the wing leading edge (Figure 1A), generating continuous airflow across the wing surface. Localised sinusoidal airflow oscillations were delivered through a ∼2mm diameter tube attached to a small electronic audio speaker driven MATLAB-generated voltage outputs from a NI-DAQ (National Instruments, USA). The tube was positioned either at the forewing base or the distal forewing (located at ∼75% spanwise from the base and 50% chordwise). Localised airflow was delivered at a range of frequencies between 4-400 Hz for 1 second per frequency. Airflow pressure exiting the tube was calibrated to equal amplitudes across the stimulation frequency range using a pressure sensor.

Wing deformation was recorded using a single high-speed camera (FLIR BFS-U3-04S2M-CS, configured to 868.8 fps), positioned to view the animals right fore/hindwing from a lateral viewing angle. The camera was synchronised with electrophysiology via an LED trigger controlled by a Teensy microcontroller; LED flashes were visible in the recorded and simultaneously logged in the electrophysiology data.

### Data Processing

During the electrode placement, we attempt to isolate units to minimize the need for spike sorting. For post-processing, neural units were detected using a single voltage threshold and manually sorted by spike waveform principal components (PCs) using Spike2 (CED, UK). Units were included for further analysis if they possessed a single distinctive spike waveform PC cluster that was stable throughout the recording session, in addition to minimal inter-spike interval refractory violations under 2 ms. Each recording (electrode penetration) typically yielded one single high-quality neural unit. For each animal, only one instance of a given neuron type was included in the analysis (e.g. if two DNs recorded in separate penetrations were both classified as WFDN_L_, only one instance was included).

Local preferred direction tuning curves were determined from average spike rates of 2 second sequential presentations of gratings moving along eight different directions (Supple et al., 2026). Temporal frequency tuning curves were calculated from average spike rates for 5 second presentations of gratings moving along the preferred direction for temporal frequencies ranging from 0.1 to 60 cycles/sec. Spike triggered averages (STAs) of gratings velocity were calculated by averaging grating velocity within a -100 ms to +50 ms window surrounding each spike time. Neuron latency was calculated from the temporal offset of maximum grating velocity preceding spike time.

Mechano-sensitivity tuning curves were calculated from spike rates in response to sinusoidally-modulated localised airflow stimuli, positioned at either the distal wing or wing base. Wing deformation (displacement) to bulk airflow was estimated from 2D digital image correlation (DIC) using Ncorr (Blaber et al., 2015). Image sequences were first rotated and cropped into a standardised view. The wing region of interest (ROI) was outlined manually on a reference image. Wing ROIs combined fore- and hindwing and were ∼96 x 150 pixels. DIC subset radius was 15–20 pixels and subset spacing 6–8 pixels to balance spatial resolution against noise; step analysis and subset truncation were enabled to handle large displacements. 2D wing displacement time series were decomposed via principal component analysis (PCA; mean-centred) to obtain the most prominent deformation modes (Yarger et al., 2025).

For spike triggered averages (STAs) of wing deformation, each PC was z-scored per trial and averaged over a -40 ms to +20 ms window per spike time. STAs of randomised spike times were also calculated to estimate the unconditioned wing deformation prior. Fourier transforms of each PC were calculated for both distal and proximal wing regions, defined by bisecting the wing chordwise at a distance halfway along span.

### Statistical Analysis

All data are reported as mean ± SEM, unless stated otherwise. Normality was assessed using the Lilliefors test. Statistical comparisons were performed using paired t-tests (within-subject). Where multiple comparisons were conducted, the familywise error rate was controlled using Bonferroni correction.

## Acknowledgements

This work was financially supported by the European Research Council (ERC-StG no.804315 ‘Vision-In-Flight’ to HTL), Biotechnology and Biological Sciences Research Council (grant BB/R002509/1 to HTL, and BB/X002276/1 to HGK), and the Air Force Office of Scientific Research (AFOSR; grant FA8655-23-1-7049 to HGK).

